# Static and dynamic measures of human brain connectivity predict complementary aspects of human cognitive performance

**DOI:** 10.1101/224956

**Authors:** Aurora I. Ramos-Nuñez, Simon Fischer-Baum, Randi Martin, Qiuhai Yue, Fengdan Ye, Michael W. Deem

## Abstract

In cognitive network neuroscience, the connectivity and community structure of the brain network is related to cognition. Much of this research has focused on two measures of connectivity – modularity and flexibility – which frequently have been examined in isolation. By using resting state fMRI data from 52 young adults, we investigate the relationship between modularity, flexibility and performance on cognitive tasks. We show that flexibility and modularity are highly negatively correlated. However, we also demonstrate that flexibility and modularity make unique contributions to explain task performance, with modularity predicting performance for simple tasks and flexibility predicting performance on complex tasks that require cognitive control and executive functioning. The theory and results presented here allow for stronger links between measures of brain network connectivity and cognitive processes.

## 1. Introduction

Research in cognitive neuroscience has typically focused on identifying the function of individual brain regions. Recent advances, however, have led to thinking about the brain as consisting of interacting subnetworks that can be identified by examining connectivity across the whole brain. This emerging discipline of cognitive network neuroscience has been made possible by combining methods from functional neuroimaging and network science (Bullmore et al., 2009; Medaglia et al., 2015; Sporns, 2014). Functional neuroimaging methods provide a rich source of data for characterizing the connections – either functionally or structurally – between different brain regions. Using these data, network science provides mathematical tools for investigating the structure of the brain network, with brain regions serving as nodes, and the connections between brain regions serving as edges in the analysis.

Under this framework, the structure of the brain network can be characterized with a variety of measures. For example, one measure, network modularity, characterizes the extent to which a network has community structure by dividing the brain into different modules so as to maximize the relative strength of within-module to between-module connections (Newman, 2006). A different measure, network flexibility, characterizes how frequently regions of the brain switch allegiance from one module to another, over time (Bassett et al., 2010). Going forward, a major challenge of cognitive network neuroscience is to determine the relationship between measures of brain network structure and cognitive theorizing (Sporns, 2014).

Relating individual differences in brain network structure to behavioral performance on cognitive tasks provides one tool for understanding the link between network measures and cognitive theories. Both modularity and flexibility have been shown to explain variation in cognitive performance. Individual differences in modularity correlate with variation in memory capacity (e.g. Meunier et al., 2014; Stevens et al., 2012)and have been shown to have a systematic relationship with task complexity, with high modular individuals performing better on simpler tasks and low modular individuals performing better on more complex tasks (Yue et al., submitted) consistent with theoretical work on modularity drawn from theoretical biology (Deem, 2013). Variation in network flexibility has been linked to other aspects of cognition including skill learning (e.g. Bassett et al., 2013, 2010), cognitive control (e.g. Alavash et al., 2015; Braun et al., 2015), and mood (Betzel et al., 2016), and has been identified as a biomarker of the cognitive construct of cognitive flexibility (Braun et al., 2015).

On the surface, this prior literature suggests that network modularity and flexibility may be tapping into different cognitive capacities. There is an intuitive basis for this distinction, as flexibility relates to how much brain networks change over time, and modularity relates to differences in interconnectivity. However, such a conclusion would be premature, since each of these previous studies measured modularity and flexibility in isolation, without considering whether the other measure could also explain variation in cognitive performance. Indeed, no study has addressed the basic question of the relationship between these two common measures of network structure, either from an empirical or theoretical perspective. This relationship might be the key to understand how different measures from network neuroscience relate to different cognitive functions.

The current study investigated the relationship between flexibility and modularity, demonstrating a strong relationship between the two measures and presenting a theoretical framework that explains this relationship. Despite this correlation, we suggest that modularity and flexibility still reflect distinct cognitive abilities, as flexibility was found to be a better predictor of performance for complex tasks while modularity was a better predictor of performance for simple tasks.

## 2. Methods

### 2.1 Participants

Participants were 52 (18-26 years old, Mean: 19.8 years; 16 males and 36 females) students from Rice University with no neurological or psychiatric disorders. Subjects were given informed consent in accordance with procedures approved by the Rice University Institutional Review Board. Subjects were compensated with $50 upon their participation in both the behavioral and imaging sessions.

### 2.2 Resting-state fMRI

#### 2.2.1 Imaging data acquisition

A high-resolution T1-weighted structural and three resting state functional scans were acquired using a 3T Siemens Magnetom Tim Trio scanner equipped with a 12-channel head coil. Scanning was done at the Core for Advanced Magnetic Resonance Imaging (CAMRI), at Baylor College of Medicine. The T1-weighted structural scan was collected prior to the functional scans and it involved the following parameters: TR=2500ms, TE=4.71ms, FoV=256mm, voxel size = 1x1x1 mm^3^. Functional runs were three 7-minute resting-state scans obtained by using the following sequences: TR = 2000ms, TE = 40ms, FoV = 220 mm, voxel size = 3x3 mm^2^, slice thickness = 4mm. A total of 210 volumes per run each with 34 slices were acquired in the axial plane to cover the whole brain. All 52 subjects participated in the imaging session.

#### 2.2.2 Preprocessing

Image preprocessing was conducted using the AFNI_2011_12_21_1014 version software (Cox, 1996). The first 6 volumes of each functional run were discarded to allow stabilization of the BOLD signal. Each functional run was preprocessed separately, including de-spiking of large fluctuations for some time points, slice timing and head motion correction. Then each subject’s functional images were aligned to that individual’s structural image, warped to the Talairach standard space, and resampled to 3-mm isotropic voxels. Next, the functional images were spatially smoothed with a 4-mm full-width half-maximum Gaussian kernel. A whole brain mask was then generated and applied for all subsequent analysis. Bandpass (0.005-0.1Hz) filtering and outlier censoring were then conducted. The outliers censoring removed the time points in which the head motion exceeded a distance (Euclidean Norm) of 0.2mm respect to the previous time point, or in which > 10% of whole brain voxels were considered as outliers by AFNI’s 3dToutcount. A multiple regression model was then applied to each voxel’s time series to regress out several nuisance signals, including third-order polynomial baseline trends, six head motion correction parameters, and six derivatives of head motion. The residual time series after application of the regression model were used for the following network analyses.

#### 2.2.3 Network re-construction, modularity, and flexibility calculation

The whole brain network was re-constructed based on 84 Brodmann areas (42 Brodmann areas for left and right hemispheres respectively). First, Brodmann area masks were generated using the TT_Daemon standard AFNI atlas (Lancaster et al., 2000), from AFNI_2011_12_21_1014 version. Then the mean time series for each area was extracted by averaging the preprocessed time series across all voxels covered by the corresponding mask. In the network, each Brodmann area served as a node and the edge between any two nodes was defined by the Pearson correlation of the time series for those two nodes. For each subject and each run, edges for all pairs of nodes in the network were estimated, resulting in an 84×84 correlation matrix. An averaged correlation matrix across three runs was then obtained for each subject. This matrix was later used to calculate modularity for each subject by applying Newman’s algorithm (Newman, 2006).

Modularity is a measure of the excess probability of connections within the modules, relative to what is expected by random chance. To calculate modularity, we first took the absolute values of each correlation and set all the diagonal elements of the correlation matrix to zero. Since fewer than .05% of the elements in the matrix were negative and their absolute values are relatively small, taking absolute values should not have major effect on results. The resulting matrix was binarized by setting the largest 400 edges in the network to 1 and all others to 0. To show that the results were persistent with the cutoff, we also considered 300 and 500 edges. This binarization process has been argued to improve detection of modularity, by increasing the signal-to-noise ratio (Chen and Deem, 2015). Modularity was defined as in Equation 1 below, where *A_ij_* is 1 if there is an edge between Broadmann areas *i* and *j* and zero otherwise, the value of *a_i_* = Σ_j_ *A_ij_* is the degree of Brodamann area *i*, and e = ½ Σ_i_ *a_i_* is the total number of edges, here set to 300,400 and 500, respectively. Newman’s algorthim was applied to the binarized matrix to obtain the (maximal) modularity value and the corresponding partitioning of Brodmann areas into different modules for each subject.

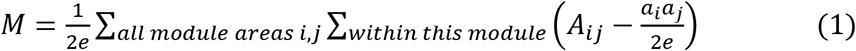

Flexibility was calculated from the time series data by computing C_i_(t), the record of which module the ith Brodmann area is in at time window t, 1≤t≤165. Flexibility for a given Brodmann area of a given subject is the number of changes in the value of Ci(t) across the 165 time windows of length 40. Results for given subject are averaged over all Brodmann areas of that subject. Finally, the average flexibility of Brodmann area i was computed by the average over all subjects of the flexibility in each subject of Brodmann area i.

#### 2.2.4 Simulations

Two simulation studies were carried out using 400 edges to determine if the relationship between flexibility and modularity observed in the human subjects could be observed with spatially correlated random data. For either study, each run of simulation generated 52 subjects (same number as the human subjects), and flexibility and modularity were calculated for each subject, thus obtaining the correlation coefficient r. Each study contains 1000 runs of simulation, and therefore corresponds to a distribution of 1000 values of r.

In both simulation studies, we generated neural activity signals with spatial structure:

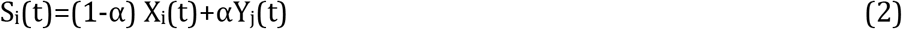

Where S_i_ denotes the signal for Brodmann area 1≤i≤84, X_i_ is a Gaussian signal different from area to area and Y_j_ is a Gaussian signal shared by areas that belong to the same module 1≤j≤4, but different from module to module. α therefore describes how modular the signal is (0≤α≤1). Specific expressions for X_i_ and Y_j_ are:

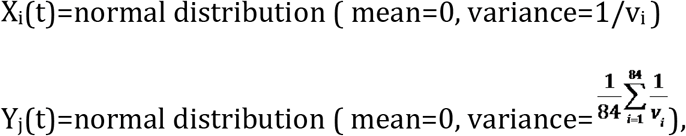

Where v_i_ is the number of voxels of area i. Note that we defined four modules in the simulation, since most of the human subjects have four modules. The module sizes were calculated as the average of the ranked values of the module sizes from the human subjects with four modules: 15, 18, 23 and 28. For each time point, X_i_ was generated for each area, and Y_j_ was generated four times corresponding to four modules. Each area’s signal at time t is then (1-α) X_i_(t)+αY_j_(t). This procedure was repeated 204 times to generate a 204*1 time series for each area of each subjects, as the real subjects also have 204 time points in their data.

The modularity of the matrix increases monotonically with α, where α=0 gives uncorrelated data. In the first simulation study (simulation 1), we chose a fixed α so that the mean value of modularity is the value from human data (M=0.48). In a second simulation study (simulation 2), to better represent the distribution of human modularity values, we varied α for each computational subject, choosing the value that produced modularity matching the corresponding human subject.

### 2.3 Behavioral tasks

All fifty-two subjects participated in the operation span and task-shifting tasks. Forty-three of them participated in the ANT task and visual short term memory task, and forty-four subjects participated in the traffic light task and digit span task, as these were done in a different session, and not all subjects returned to participate in all tasks. The interval between neuroimaging and behavioral sessions varied from 0 (i.e., measuring resting-state fMRI and behavior on the same day but during different sessions) to 140 days.

#### 2.3.1 Operation span

Subjects were administered the operation span task (Unsworth et al., 2005) to measure their working memory capacity. This task has been shown to have high test-retest reliability, thus providing a stable measure in term of the rankings of individuals across test session (Redick et al., 2012). In this task, for each trial, participants saw an arithmetic problem, e.g., (2×3)+1, and were instructed to solve the arithmetic problem as quickly and accurately as possible. The problem was presented for 2 seconds. Then, a digit, e.g., 7, was presented on the next screen. Subjects judged whether this digit was a correct solution to the previous arithmetic problem by using a mouse to click a “True” or “False” box on the screen. After the arithmetic problem, a letter was presented on the screen for 800ms that subjects were instructed to remember. Then the second arithmetic problem was presented, followed by the digit and then the second letter, with the same processing requirements for both arithmetic problem and letter, and so forth. At the end of each trial, subjects were asked to recall the letters in the same order in which they were presented. The recall screen consisted of a 3×4 matrix of letters on the screen and subjects checked the boxes aside letters to recall. Subjects used the mouse to respond to the arithmetic problem and to recall letters. The experimental trials included set sizes of six or seven arithmetic problem – letter pairs. There were twelve trials for each set size, resulting a total of 156 letters and 156 math problems. The six and seven set size trials were randomly presented.

Before the actual experiment, a practice session was administered to familiarize subjects with the task. The practice session consisted of a block involving only letter recall, e.g., recalling 2 or 3 letters in a trial, a block involving only arithmetic problems, and a mixed block in which the trial had the same procedure as in the experimental trials, i.e., solving the arithmetic problems while memorizing the letters, and recalling them at the end, but with smaller set sizes of 2, 3, or 4.

The response times for math problems and accuracy for arithmetic problems and letter recall were recorded. The operation span score is the accuracy for letter recall, calculated as the number of letters that were recalled at the correct position out of total number of the presented letters. The maximum span score is 156.

#### 2.3.2 Visual arrays task

A visual array task was used to tap visual short-term memory capacity. In this task, subjects were instructed to fixate at the center of the screen. Arrays of 2 to 5 colored squares at different positions on the screen were presented for 500ms, followed by a blank screen for 500ms, and then by multi-colored masks for 500ms. A single probe square was then presented at one of locations where the colored squares had appeared. Subjects had to judge whether the probe square had the same or different color as the one at the same position. The order of different array sizes was random. Each array size condition had 32 trials, half of which were positive response trials and half negative. We calculated the visual short-term memory capacity (Cowan, 2000; Rouder et al., 2011) as k=N×(H-F), where N was the number of colored squares in the largest set size (here N=5), H was the hit rate, and F was the false alarm rate.

#### 2.3.3 Digit span

In this task a list of numbers were presented in auditory form at the rate of one number per second and participants were required to memorize them. After presenting the last number in the list, a blank screen prompted participants to recall the numbers in the order in which they were presented by typing on the keyboard. Participants were given five trials for each set size starting at two. The program would terminate if participants got less than 3 correct trials for that set size (60% accuracy). Digit span was calculated by estimating the list length at which the subject would score 60% correct using linear interpolation between the two set sizes that spanned this threshold.

#### 2.3.4 Task-shifting task

In this task, participants responded to an object according to a preceding cue word. The object was either a square or a triangle, and the color of the object was either blue or yellow. If the cue was “color”, participants pressed a button to indicate whether the object was blue or yellow, and if the cue was “shape”, they pressed a button to indicate whether the object was a square or a triangle. The same buttons were used for the two tasks. The response time was recorded from the onset of the object. For half of the trials, the cue was the same as that in the previous trial, a repeat trial, and for the other half, the cue changed, a switch trial. For each condition, to take into account both response time and accuracy in a single measure, we calculated the inverse efficiency (IE) score (Townsend and Ashby, 1983) defined as mean RT/proportion correct. The task shifting cost was measured as the difference in inverse efficiency score between the repeat and switch trials. We also adjusted cue-stimulus Interval (CSI), which is the time between onset of the cue and onset of the object, using CSIs of 200ms, 400ms, 600ms, and 800ms. However, as the effect of modularity on IE did not differ for different CSIs, the data were averaged across CSI. In total, there were 256 repeat trials and 256 switch trials.

#### 2.3.5 Attention network test

The Attention Network Test (ANT; Fan et al., 2002) was used to measure three different attentional components: alerting, orienting, and conflict resolution. In this task, subjects responded to the direction of a central arrow, pressing the left or right mouse button to indicate whether it was pointing left or right. The arrow(s) appeared above or below a fixation cross, which was in the center of the screen. The central arrow appeared alone on a third of the trials and was flanked by two arrows on the left and two on the right on the remaining two third of trials. The flanking arrows were evenly split between a condition in which they pointed in the same direction as the central one, a congruent condition, and a condition in which they pointed in the opposite direction, an incongruent condition. In the neutral condition, there were no flanking arrows. On three-quarters of the trials, the arrow(s) were cued by an asterisk or two asterisks, which appeared for 100ms on the screen. The interval between offset of the cue and onset of the arrow was 400ms. There were four cue conditions: 1) no cue condition, 2), a cue at fixation, 3) double-cue condition, with one cue above and the other below fixation, 4) spatial-cue condition, where the cue appeared above or below the fixation to indicate where the arrows would appear. Thus, the task had a 4 cue × 3 flanker condition factorial design. The experimental trials consisted of three sessions, with 96 trials in each session, and 8 trials for each condition. For half of all trials, arrows were presented above the fixation and for the other half below. Also, for half of the trials, the middle arrow pointed left and for the other half right. The order of trials in each session was random. Before the experimental trials, 24 practice trials with feedback were given to subjects that included trials of all types.

Response times and accuracy were recorded. Mean RT for each condition for each subject was computed based on correct trials only. As with task shifting, we calculated the inverse efficiency (IE) score for each condition. The alerting effect was computed by subtracting the IE for the no cue condition from the IE for the double cue condition. The orienting effect was computed by subtracting the IE for the center cue condition from the IE for the spatial cue condition. The conflict effect was computed by subtracting the IE for the congruent condition from the IE for the incongruent condition. To make the direction of the conflict effect the same as that of alerting and orienting effects, we reversed the sign of conflict effect. Thus, the more negative the conflict effect value, the greater the interference from the incongruent flankers.

#### 2.3.6 Traffic light task

In this task, subjects saw a red square in the center of screen, which was replaced after an unpredictable time delay (from 2 to 3 seconds) by a green circle. Subjects pressed a button as quickly as possible when they saw the green circle. There were 25 trials in total. Mean response time was calculated for each subject.

#### 2.3.7 Simple vs. Complex Composite scores

Previous empirical research (Yue et al., submitted) and theoretical work (Deem, 2013) suggests an interaction between measures of network structure and performance on simple versus complex tasks. For the purposes of the current research, complex tasks are defined as those whose performance taps into executive functions and cognitive control, while simple tasks do not depend on these operations. From the battery of tasks described above, complex tasks include operation span, visual arrays, digit span, task-shifting, and the conflict resolution component of the ANT, while simple tasks include the orienting and alerting components of the ANT and the traffic light task. To limit the number of behavioral measures in the analysis, two composite scores, simple and complex were calculated for the 40 subjects who participated in all of the tasks described above. These composites were computed by summing the z-scores for the performance measures for the simple and complex tasks.

## 3. Results

### 3.1 Correlations of modularity and flexibility

Using Brodmann areas as nodes and functional connectivity between these nodes (determined from resting state fMRI) as the measure of the strength of edges, we determined modularity and flexibility for each subject. Figure 1a depicts the relationship between modularity and flexibility across our 52 participants. For the 400-edge analysis, modularity values ranged from .33 to .60, with a mean of .48 (standard deviation 0.055) on a scale from 0 to 1.0. Flexibility values ranged from 26.4 to 38.7 with a mean of 31.6 (standard deviation 2.79). A striking negative correlation r=-0.78 (p < 0. 000001) was obtained between these two mathematically different measures, which had not been previously reported. The analysis for the 300- and 500-edge yielded negative correlations of r=-0.70 (p < 0.000) and r=-0.74 (p < 0.000) respectively.

To understand the extent to which modularity and flexibility are necessarily related, two simulation studies were carried out (see Figure 1b). In simulation 1, connectivity data were generated such that the mean modularity across different connectivity matrices matched the mean for the human data, but the standard deviation of the modularity values was significantly lower than the standard deviation of the actual population. The distribution of r-values (blue color bars in Figure 1b) has a mean of -0.16, with all thousand r-values below the negative correlation of -.78 observed in the human data (p < .001). It is worth noting that the standard deviation of the modularity values is narrow in this stimulation (0.028), and smaller than that of the human data. The second simulation matched the human data on individual modularity values, thus matching both their mean and standard deviation. As shown in Figure 1b (red color bars), this model results in correlation coefficients consistent with the value measured for human subjects, with an average -0.72 and a standard deviation 0.05 (p > .2). Thus, it is the diversity in human modularity values that leads to the negative correlation between flexibility and modularity.

### 3.2 Consistency of constituent BAs in modules and flexibility across BAs

As reported in Yue et al. (submitted), the most typical number of modules for each subject was 4, with a range of 3 to 6. The four modules roughly corresponded to networks for 1) somatosensory-motor processing, 2) bilateral auditory/language processing, 3) default mode processes, and 4) a diverse set of functions including visual processing, attention, and memory. In Yue et al., (submitted) we quantified the degree of consistency in assignment of BAs to modules across subjects by a distance measure which was a count of the number of subjects for whom the assignment of a BA to module for their own data differed from the assignment based on group average data. For low-level sensory-motor BAs, the assignment of nodes to modules was highly consistent across subjects whereas for high-level cognitive control areas, the assignment was highly variable.

We also analyzed the flexibility of each Brodmann area. Some BAs more frequently changed module alignment across time, and therefore had higher flexibility scores, than others. Figure 1c illustrates flexibility by BA. Regions with higher flexibility include the anterior cingulate cortex, ventromedial prefrontal cortex, orbitofrontal cortex, and dorsolateral prefrontal cortex bilaterally, regions typically associated with cognitive control and executive functions. Regions with lower flexibility are those regions involved in motor, gustatory, visual, and auditory processes such as postcentral gyrus, primary motor cortex, primary gustatory cortex, and secondary visual cortex. In general, the regions showing greater distance in module assignment were the same regions showing greater flexibility, with a strong correlation between distance and flexibility across BAs (r=.64, p=.001).

### 3.3 Relationship with cognitive performance

Results for the seven behavioral tasks are reported in supplementary materials and in Figure 1 supplemental. Scores from the seven behavioral tasks were converted into z-scores and combined to create simple and complex composite scores. Correlation analyses revealed a significant negative correlation between modularity measured with 400 edges and the complex composite (r=-0.330, p=0.038). In contrast, for simpler tasks, individuals with high modularity performed better, with a significant positive correlation between modularity and simple composite (r=0.449, p=0.004). As might be expected, given the strong negative correlation between modularity and flexibility, there was a significant positive correlation between flexibility measured with 400 edges and the complex composite (r=0.408, p=0.009) and a negative correlation with the simple composite (r=-0.242, p=0.133). The same pattern was observed at different edge densities (see Table 1).

Despite the strong correlation between flexibility and modularity, it is possible that they make independent contributions to explaining individual differences in cognitive performance. As show in Figure 2 and Table 1, the magnitude of the correlation coefficient between modularity and the simple task composite is larger than the correlation coefficient between flexibility and simple task composite, across edge densities. The opposite pattern is true for the complex tasks. The correlation coefficient between flexibility and task performance is higher than the correlation coefficient between modularity and task performance. This pattern is confirmed by multiple regression analysis performed on tasks performance, regressing the effect on modularity and flexibility measured in a network with 400 edges simultaneously in order to determine the significance of the unique contribution of each. For the simple task composite, the coefficient for modularity was significant (β=.75, p<.004), but that for flexibility was not (β=.37, p<.14). In contrast, for the complex task composite, the coefficient for flexibility was marginally significant (β=.48, p<.07) but that for modularity was not (β=.09, p=.72).

**Table 1.** The correlation coefficients for simple and complex tasks at different edge densities

|  | Correlation Coefficient |  |  |  |  |  |
| --- | --- | --- | --- | --- | --- | --- |
|  | Modularity |  |  | Flexibility |  |  |
| Number of Edges | 300 | 400 | 500 | 300 | 400 | 500 |
| Simple Tasks | .43 | .45 | .39 | -.27 | -.24 | -.14 |
| Complex Tasks | -.38 | -.30 | -.32 | .40 | .41 | .33 |

## 4. Discussion

The present results show the two measures of brain network structure that are treated as independent in the literature – flexibility and modularity – are actually highly related. Still, each makes independent contributions to cognitive performance, with modularity contributing more to performance on simple tasks and flexibility contributing more to performance on complex tasks. Simulations revealed that this correlation was dependent on the distribution in modularity observed across subjects. This finding raises the question of why the population has such a broad distribution of modularity values. One explanation is that broad distribution ensures that some fraction of the population will have low M, suitable for performance on complex tasks, and some fraction will have high M, for performance on simple tasks (Deem, 2013). From an evolutionary point of view, and in light of Figure 2, selection for performance on a broad spectrum of tasks should lead to such a broad range of modularity values.

How might we account for this strong negative correlation between flexibility and modularity? An intuitive explanation for the negative correlation between flexibility and modularity derives from a dynamical systems perspective that views different configurations of brain regions as attractor states, with modularity measuring the depth of the attractor states (Smolensky et al., 1996). Flexibility measures how frequently the brain transitions between states. Deeper states will naturally be more stable and resistant to transitions, leading to a negative correlation between modularity and flexibility. Given the high degree of correlation between these two measures, it is difficult to interpret findings in the literature that report only one of these measures in isolation.

Still, flexibility and modularity make independent contributions to explaining task performance and likely link to different cognitive processes. Our results suggest that flexibility, rather than modularity, may reflect cognitive control processes (Bassett et al., 2013, 2010). The regions that show the highest flexibility (Figure 1b) are those that have been previously implicated in control. The complex tasks used in the current study all require control, while the simple tasks do not. Assuming flexibility indexes cognitive control capacity, we can explain why variation in flexibility plays a larger role in explaining performance on complex tasks.

Flexibility is only weakly related to performance on simple tasks. Why then, has flexibility been shown to have strong relationships with the ability to learn even in very simple motor tasks (Bassett et al., 2010)? We argue that at the initial stages of learning the acquisition of even simple skill benefits from cognitive control operations, making tasks appear more complex. Therefore, at the initial stages of learning, it is beneficial to have a more flexible brain (Bassett et al., 2013). As learning progresses and the task becomes automatized, cognitive control is no longer necessary and the task becomes simpler. Following the theory depicted in Figure 2, as learning progresses, flexibility should decrease and modularity should increase, as has been previously observed (Bassett et al., 2015, 2013).

## 5. Conclusion

For cognitive network neuroscience to advance, better links between measures of network structure to cognitive and neural computations must be developed (Sporns, 2014). The theory and results presented here, disentangling the effects of two commonly, but interrelated measures, are one step. By considering how different measures of brain structure relate to each other and relate to variation in performance, we can start to develop stronger links between the cognitive and the network sides of this new approach.

## Acknowledgements

We are grateful for the students who participated in this study. This work was partially supported by the T.L.L. Temple Foundation. MWD and FDY were partially supported by the Center for Theoretical Biological Physics under NSF grant #PHY-1427654.

## Supplementary Material Static and dynamic measures of human brain connectivity predict complementary aspects of human cognitive performance

### 1. Network re-construction

To demonstrate that the correlation between modularity and flexibility persists after using different functional and anatomical parcellations of the brain, the whole brain network was re-constructed based on methods used by others in the resting state literature (Craddock et al., 2012; Glasser et al., 2016; Gordon et al., 2016; Power et al., 2011). As shown in table 1, all the different parcellation methods yielded a negative correlation between modularity and flexibility. However, the magnitude of the coefficient was much larger using the Brodmann areas as the parcellation anatomical map.

### 2. Relationship with task performance

Figure 1 supplemental illustrates the relationship between modularity and task performance and flexibility and task performance during tasks varying from simple to complex processes. Simple tasks include the traffic light and the orienting effect from the Attention Network Task (ANT). Complex tasks involve the Operation Span, Digit Span, Visual Short-term Memory, the Conflict Effect from the ANT, and the Task-Shifting. For the purposes of the current research, this simple vs. complex task distinction is operationalized in the following way: complex tasks are those tasks in which executive attention and cognitive control (the ability to ignore preponderant distractors while performing correctly the task at hand) are required to properly perform the task. Simple tasks are those tasks whose performance does not depend on these operations. Because of the engagement of cognitive control, complex tasks typically require longer processing times than simple tasks. In terms of complex tasks, flexibility generally shows a larger coefficient than modularity. The opposite is true for simple tasks and modularity: modularity presents with a larger coefficient than flexibility.

### 3. Supplementary Figure captions and Tables

**Figure 1 Supplemental.**
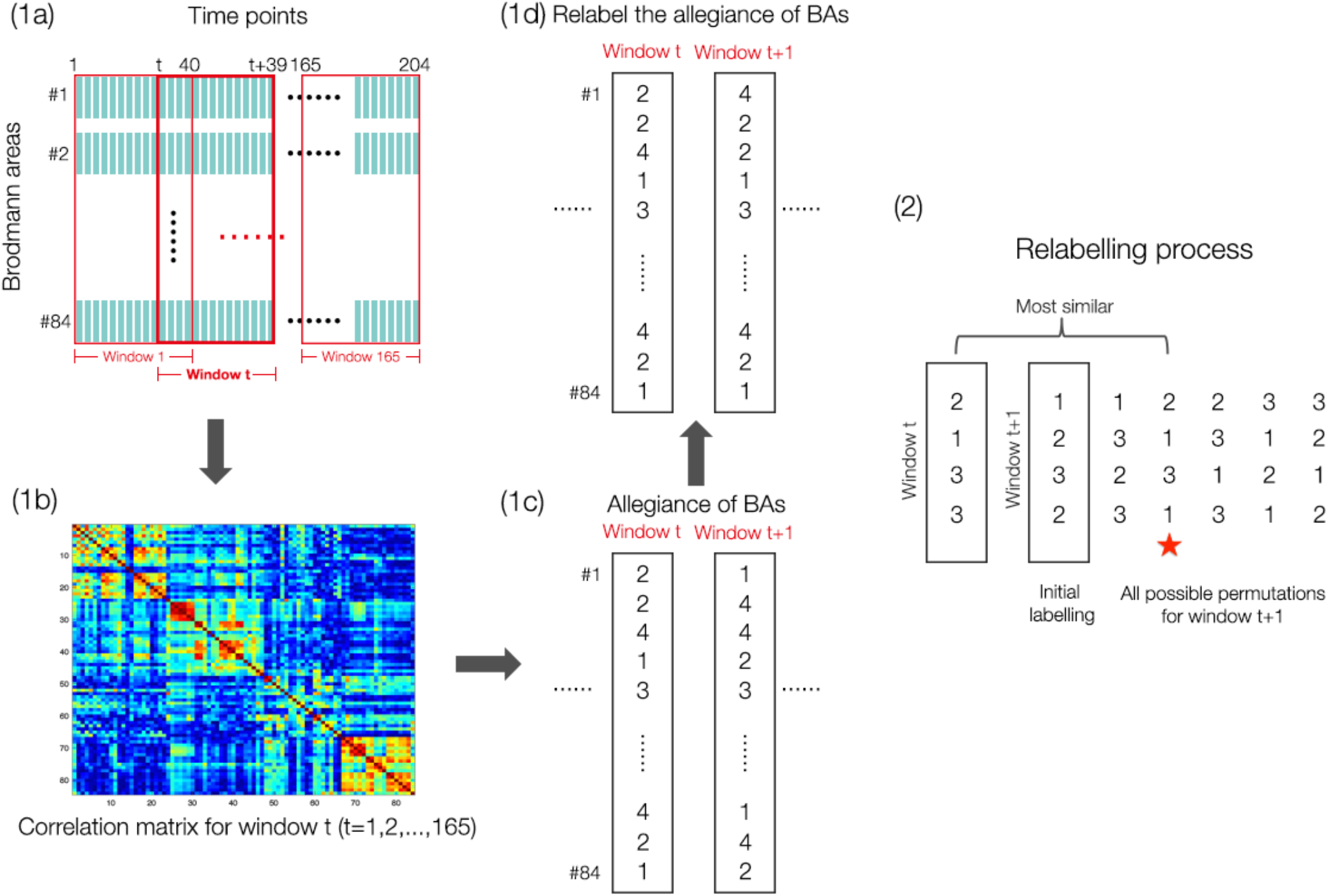
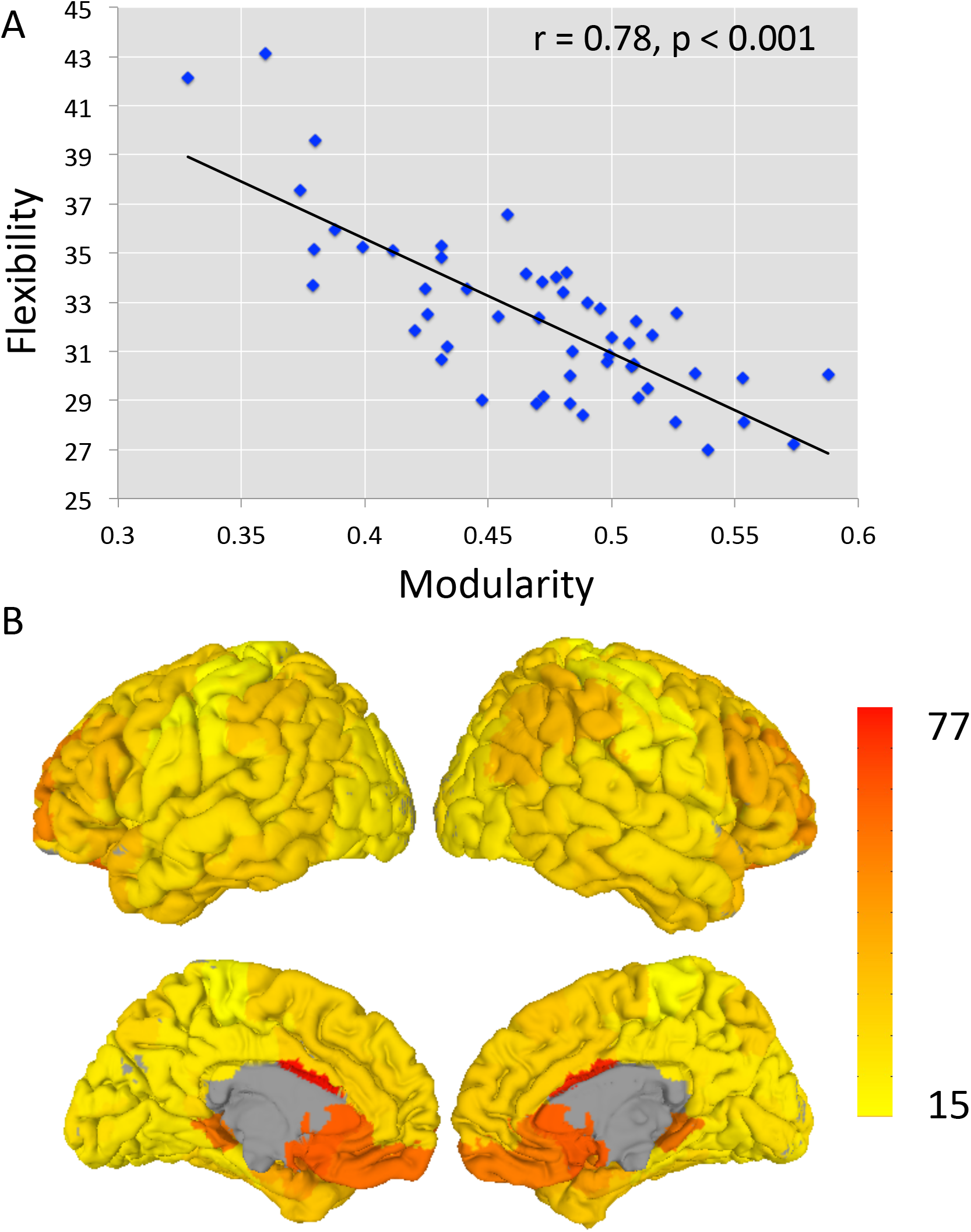
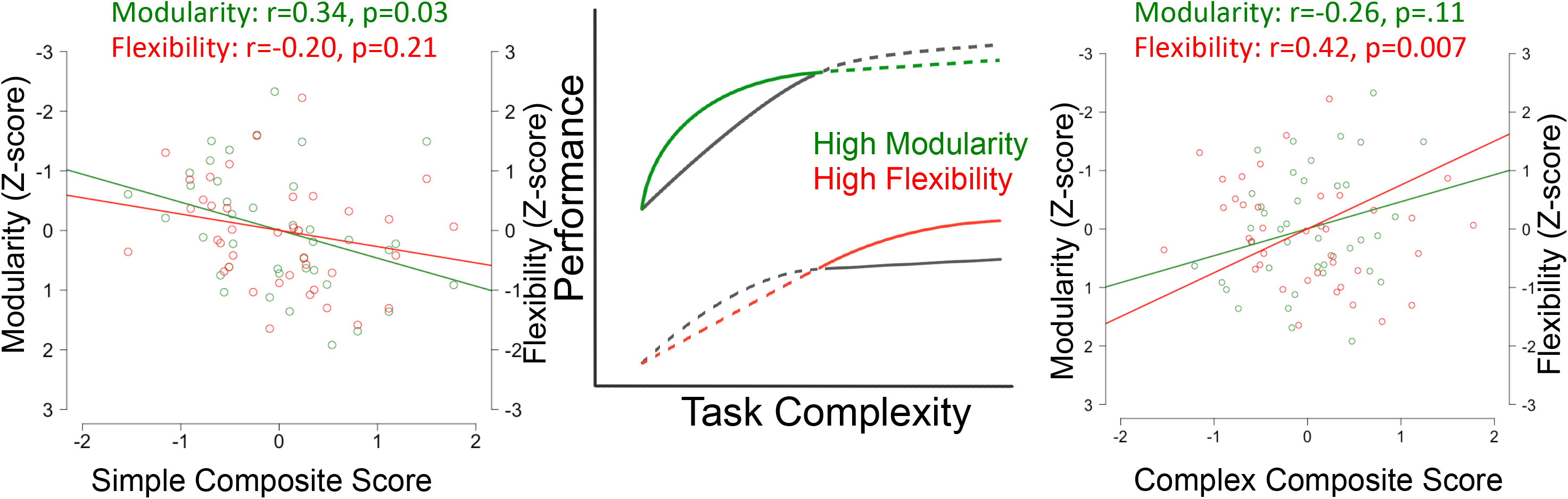
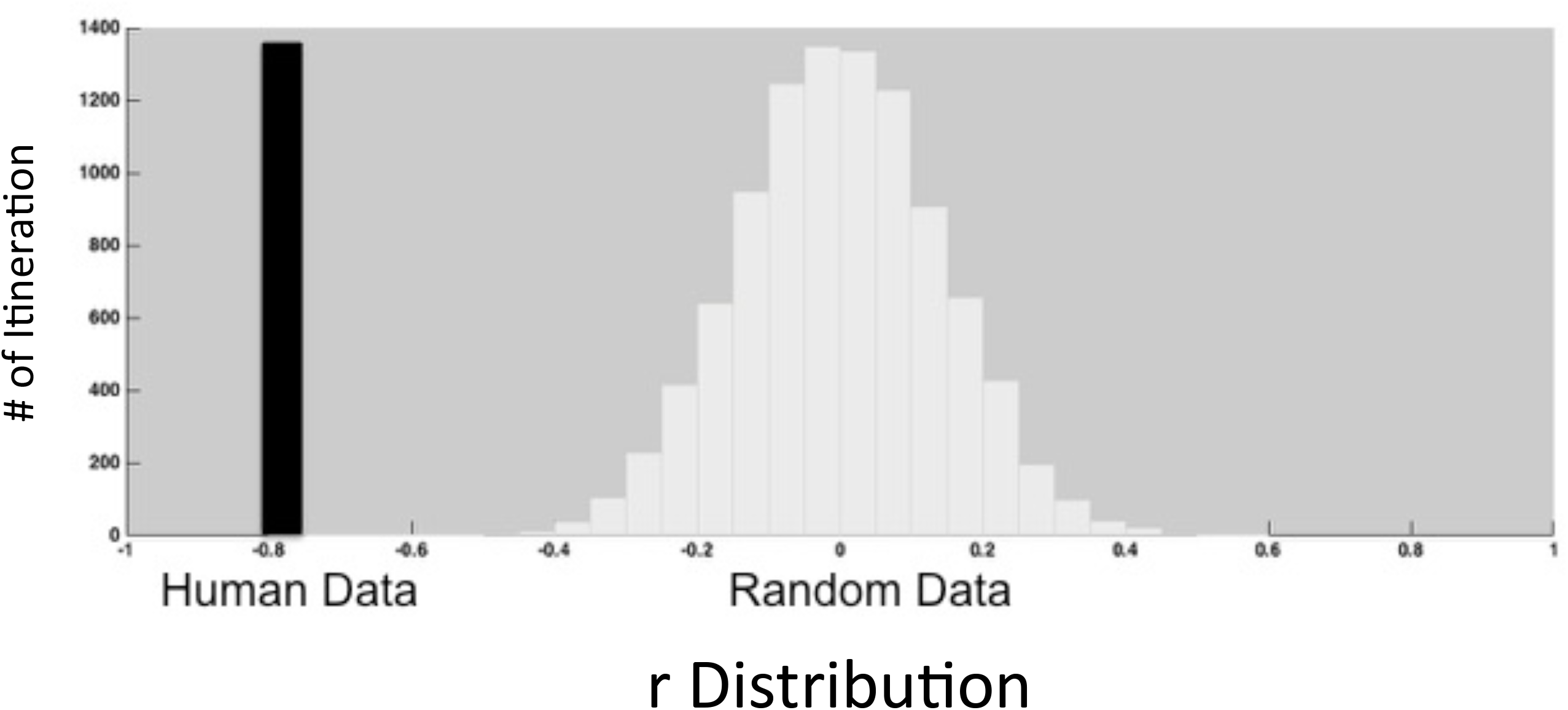
The relationship between modularity and flexibility with task performance represented by the magnitude of the coefficient between modularity and task performance and flexibility and task performance organized from simple (left) to complex (right). The center of the figure depicts the theoretical prediction relating performance to tasks at different levels of complexity for individuals with high and low modularity (green curve) and flexibility (red curve).

**Figure 2 Supplemental.**
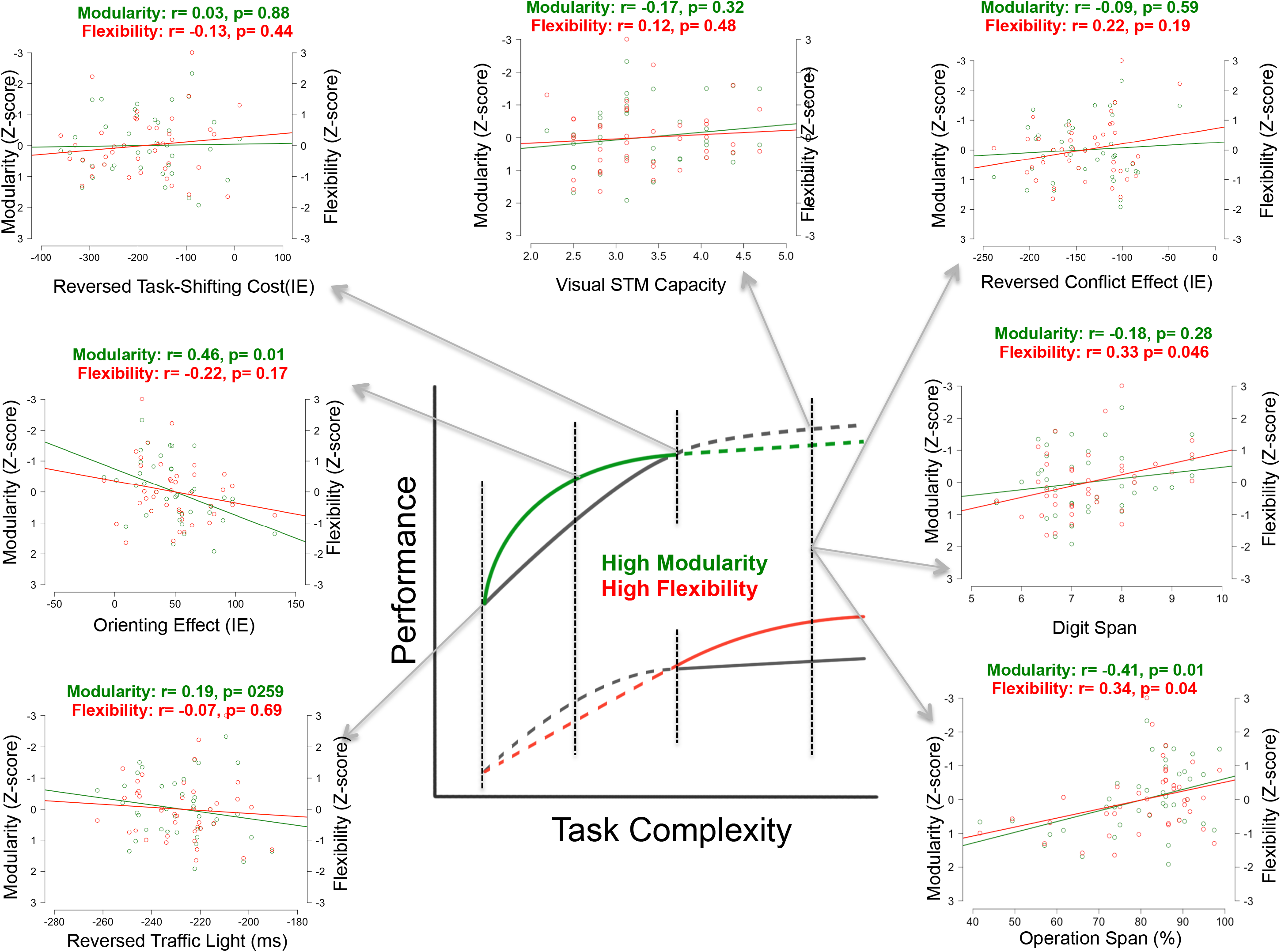
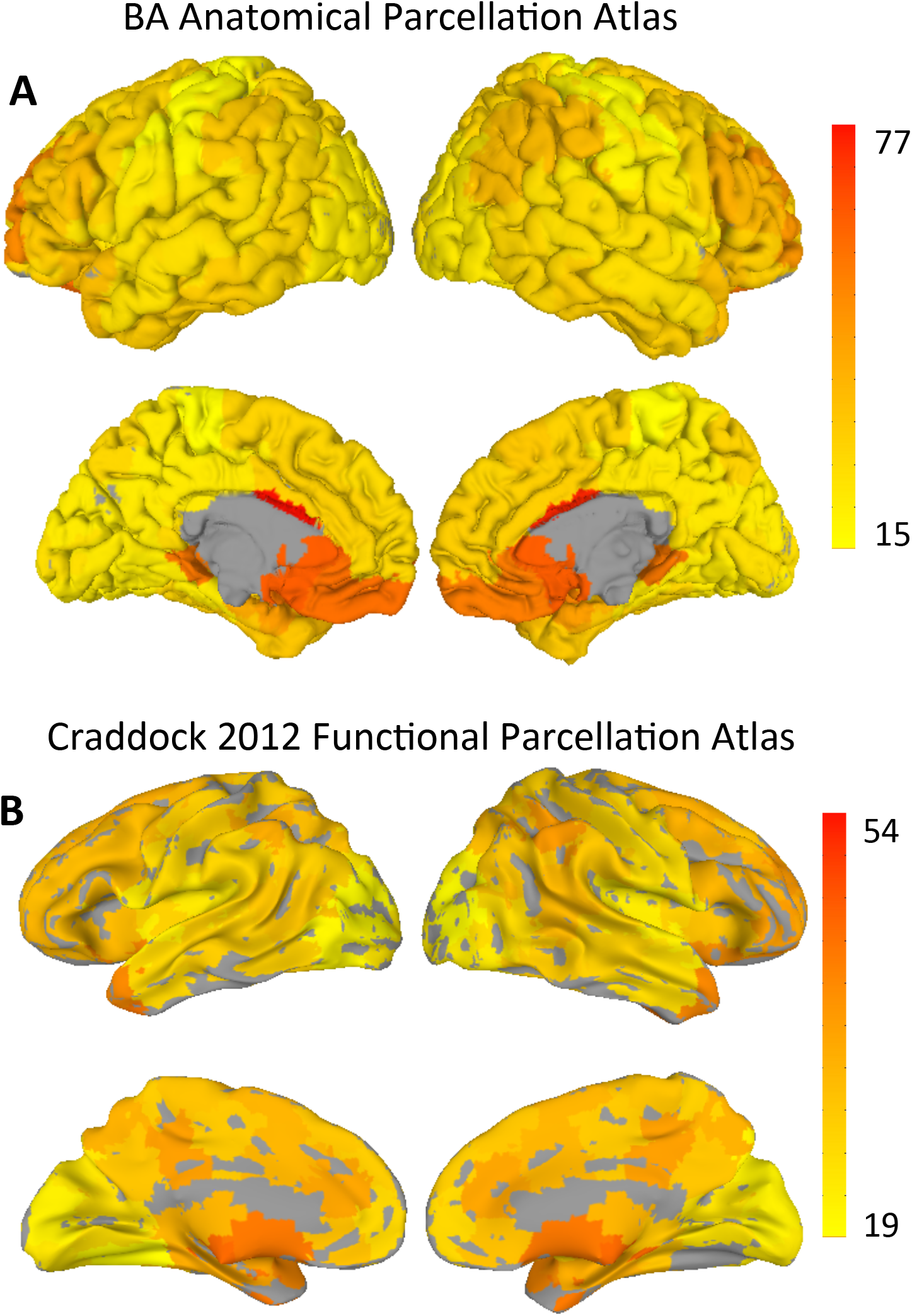
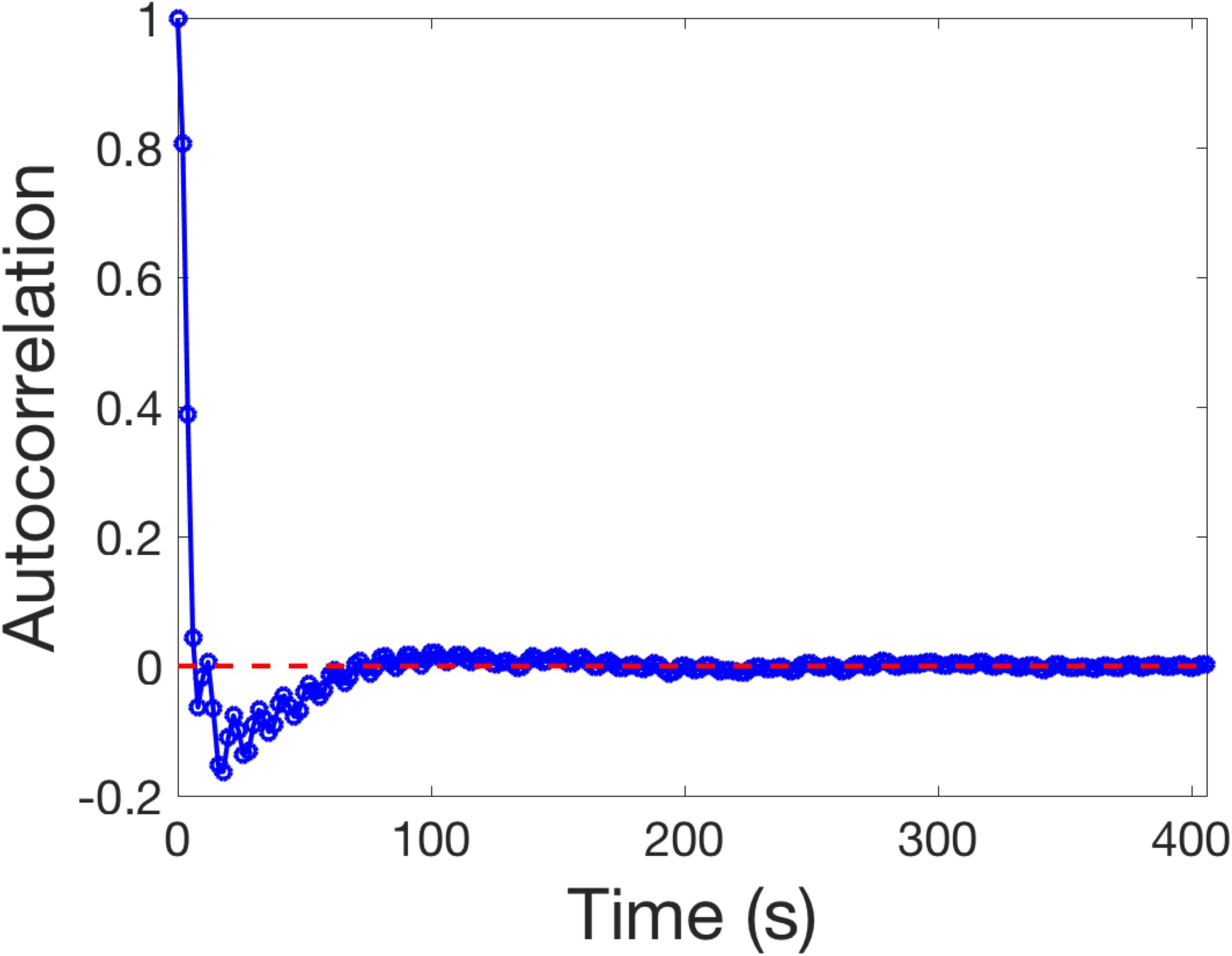
A comparison of flexibility across the brain between: A) an anatomical based parcellated atlas with 84 regions such as Brodmann’s Areas (BA) and B) a fuctional parcellated atlas with 100 regions from craddock et. al., 2012. The color bar on the right side of the figures represents flexibility values going from low (15 for BA atlas and 19 for Craddock atlas) to high (77 fir BA atlas and 54 for Craddock atlas).

**Supplementary Table 1.** Modularity and Flexibility correlation coefficients from various anatomical and functional atlases

|  | r | p-value |
| --- | --- | --- |
| BA_300_edge | -0.81 | p < 0.001 |
| BA_400_edge | -0.78 | p < 0.001 |
| BA_500_edge | -0.74 | p < 0.001 |
| Glasser et al., 2016 | -0.44 | p = 0.001 |
| Gordon et al., 2016 | -0.34 | p = 0.014 |
| Power et al., 2011 | -0.38 | p = 0.005 |
| Craddock et al., 2012 |  |  |
| 100 parcellations | -0.49 | p < 0.001 |
| 200 parcellations | -0.37 | p = 0.007 |
| 300 parcellations | -0.42 | p < 0.002 |

**Supplementary Table 2.** Modularity (M), Flexibility (F), and simple and complex task performance correlation coefficients when white matter and CSF signals were regressed out. We do not believe the white mattera and CSF signals are purely noise.

|  | r | p-value |
| --- | --- | --- |
| M&F BA_400_edge | -0.70 | 0.001 |
| M BA_400_edge & simple task performance | 0.23 | 0.161 |
| M BA_400_edge & complex task performance | -0.21 | 0.196 |
| F BA_400_edge & simple task performance | -0.07 | 0.646 |
| F BA_400_edge & complex task performance | 0.24 | 0.145 |

**Supplementary Table 3.** Correlations between individual tasks and modularity and flexibility without controlling for days in between collecting behavioral measures and resting state fMRI data

|  | Modularity |  | Flexibility |  |
| --- | --- | --- | --- | --- |
|  | r | p-value | r | p-value |
| Ospan | -0.410 | 0.011 | 0.335 | 0.040 |
| Dspan | -0.180 | 0.279 | 0.326 | 0.046 |
| Conflict | -0.089 | 0.593 | 0.219 | 0.186 |
| VSTM | -0.166 | 0.319 | 0.118 | 0.482 |
| Shifting | -0.026 | 0.875 | -0.130 | 0.438 |
| Orienting | 0.462 | 0.008 | -0.219 | 0.186 |
| Traffic | -0.190 | 0.254 | -0.066 | 0.695 |

**Supplementary Table 4.**
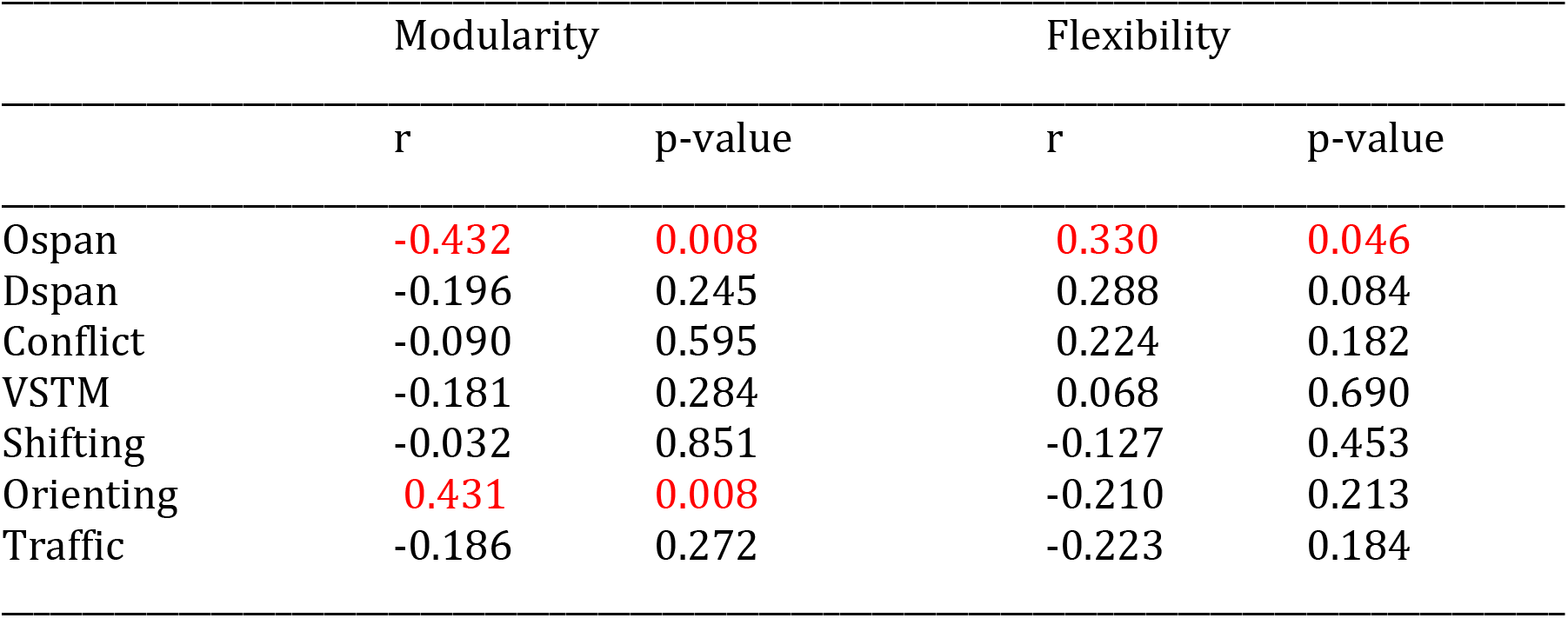
Correlations between individual tasks and modularity and flexibility controlling for number of days in between collecting behavioral measures and resting state fMRI data

